# Compatibility and Multi-Season Field Evaluation of *Trichoderma koningiopsis* Integrated with Fungicides for Soybean Charcoal Rot Management

**DOI:** 10.64898/2026.05.11.724353

**Authors:** Juliana Bleckwedel, Raúl Exequiel Nieva, Victoria González, Leonardo Daniel Ploper, Sebastian Reznikov

## Abstract

Soybean (*Glycine max* [L.] Merr.) productivity is frequently compromised by soil-borne pathogens. *Macrophomina phaseolina* (Mp), the causal agent of charcoal rot, can produce important soybean yield losses especially when hot and dry weather prevails. Integrating biological control agents with chemical seed treatments represents a promising strategy for improving disease management. This study aimed to (i) assess the *in vitro* compatibility of *Trichoderma koningiopsis* with commercial fungicide seed treatments, and (ii) evaluate the field performance of *T. koningiopsis*, alone or combined with compatible fungicides, across three soybean growing seasons. Compatibility assays revealed fungicide-specific effects, with Acronis® classified as non-fungitoxic and Topseed Extra as moderately fungitoxic. Across field seasons, Mp inoculation reduced seedling emergence, while several seed treatments improved emergence compared to the inoculated control, however, treatment effects varied markedly among years. Disease severity did not differ significantly among treatments in any season, and yield responses were strongly modified by environmental conditions rather than treatment effects. Temperature-response assays showed that *T. koningiopsis* exhibited optimal growth between 28 to 30°C and complete inhibition above 40°C, indicating high thermal sensitivity. The results demonstrate that *T. koningiopsis* can be integrated with compatible fungicides and may enhance early stand establishment under favorable conditions, but its field performance is strongly limited by high temperatures. These findings highlight the importance of environmental conditions when biological seed treatments are used.

## 1. Introduction

Soybean (*Glycine max* [L.] Merr.) is a globally significant crop and a major source of protein and oil. However, its productivity is frequently constrained by soil-borne pathogens. *Macrophomina phaseolina* (Mp), the causal agent of charcoal rot, can cause substantial yield losses, particularly under hot and dry conditions (Marquez et al., 2021). Traditional management strategies have relied predominantly on chemical fungicides, which, although effective, raise environmental concerns and may contribute to the development of pathogen resistance. Consequently, there is increasing interest in integrating biological control agents, such as *Trichoderma* species, with chemical treatments to enhance disease suppression and promote more sustainable production systems (Gordani et al. 2023; Villavicencio-Vásquez et al. 2025; Zhang et al. 2021).

*Trichoderma* spp. are well known for their antagonistic activity against diverse plant pathogens and for their capacity to stimulate plant growth. Their modes of action include mycoparasitism, competition for nutrients and space, and the induction of plant defense responses. Seed treatments with *Trichoderma* spp. have been shown to suppress pathogenic fungi and promote seedling growth in crops such as wheat (Couto et al. 2021). In soybean, *Trichoderma* spp. applied as seed treatments act as antagonists against seed- and soil-borne pathogens and as plant growth stimulators (Bleckwedel et al. 2024; Reznikov et al. 2016; Zandoná et al. 2019).

The performance of *Trichoderma*-based seed treatments is strongly influenced by environmental conditions, particularly temperature and soil moisture, which affect spore germination, root colonization, and antagonistic activity. High temperatures can reduce conidial viability, enzyme secretion, and limit rhizosphere establishment, ultimately constraining biocontrol efficacy under field conditions (Harman et al., 2021). Reznikov et al. (2016) evaluated the control of charcoal rot in soybean using *Trichoderma viride*, *Bacillus subtilis*, and chemical fungicides under field conditions. All treatments mitigated the reduction in plant emergence and disease severity caused by Mp, with the highest yields obtained in chemical treatments, followed by those treated with *Trichoderma* and *Bacillus*.

The antagonistic effect of *T. koningiopsis* against Mp has been previously demonstrated *in vitro*, under greenhouse conditions, and in field trials with artificial inoculations across two soybean seasons (Bleckwedel et al., 2024). These studies showed that *T. koningiopsis* had a biocontrol effect against Mp under greenhouse and field conditions. Similarly, it improved the weight and length of soybean plants under greenhouse conditions.

Although previous studies have demonstrated the antagonistic potential of *T. koningiopsis* against *M. phaseolina*, little is known about its thermal tolerance and its capacity to persist under the high-temperature regimes typical of charcoal rot environments. Thermal sensitivity varies widely among *Trichoderma* species and even among isolates, affecting their ecological fitness and field performance (Adhikari, P., 2023; Di Lelio et al., 2021). Therefore, characterizing the temperature–growth relationship of *T. koningiopsis* is essential to understand its limitations and to interpret its field behavior across contrasting seasons.

The integration of *Trichoderma* spp. with chemical fungicides offers a promising approach to disease management. This combined strategy can enhance the efficacy of pathogen control while potentially reducing the quantity of chemical inputs required. Zandoná et al. (2019) demonstrated that the combination of fludioxonil + *Trichoderma* spp. as soybean seeds treatment under field condition increased crop development and grain yield respect to the control.

Despite these promising findings, the effectiveness of combined *Trichoderma*–fungicide seed treatments can vary substantially depending on environmental conditions and pathogen pressure. Further investigation is required to optimize treatment protocols and to understand the interactions between biological and chemical agents under diverse field conditions. For that reason, the present study aimed to (i) assess the *in vitro* compatibility of *Trichoderma koningiopsis* with commercial fungicide seed treatments, (ii) evaluate the field performance of *T. koningiopsis*, alone and combined with compatible fungicides, across three soybean growing seasons, and (iii) evaluate the thermal sensitivity of *T. koningiopsis*.

The hypotheses of this study are that (i) *T. koningiopsis* will remain viable when combined with non-fungitoxic seed treatments, (ii) compatible combinations will improve stand establishment in field trials, and (iii) high temperatures will limit the field performance of the biocontrol agent.

## 2. Materials and methods

### 2.1 Fungal isolates and plant material

#### Thichoderma koningiopsis

The *T. koningiopsis* isolate used in this study was previously obtained and characterized by Bleckwedel et al. (2024). The isolate is preserved at −20°C on filter paper and was cultured on acidified potato dextrose agar medium (APDA, pH 6) 7 days prior to each assay.

#### Macrophomina phaseolina

The Mp17 isolate, originally collected from symptomatic soybean plants in northwestern Argentina (Reznikov et al., 2018), was used for all inoculations. The isolate is preserved at −20°C on filter paper and was cultured on acidified potato dextrose agar medium (APDA, pH 6) 5 days prior to each assay.

#### Soybean cultivars

Charcoal rot–susceptible soybean cultivars from maturity group VI were used: IS 62.1 (2021/2022), RA 655 (2023/2024), and DM 60i62 (2024/2025). Seeds were obtained from certified commercial lots.

### 2.2 *In vitro* compatibility of *T. koningiopsis* with chemical seed treatment fungicides

Eight commercial fungicide formulations commonly used for soybean seed treatment were evaluated (Table 1). Stock solutions (1000 μg/mL a.i.) were prepared in distilled water and incorporated into PDA to obtain final concentrations of 0, 0.1, 1, 10, 100 and 1000 ppm.

**Table 1.** Commercial fungicides, active ingredients and chemical group used for *in vitro* compatibility test of *T. koningiopsis*.

| Commercial Fungicide | Active Ingredient | Chemical Group |
| --- | --- | --- |
| **Apron® Maxx RFC** | metalaxyl-M 3.75% | phenylamide |
|  | fludioxonil 2.50% | phenylpyrrole |
| **Cruiser® Plus** | fludioxonil 2.50% | phenylpyrrole |
|  | difenoconazole 2.50% | triazole |
|  | thiamethoxam 26.20% | insecticide |
| **Cruiser® Advance** | fludioxonil 2.50% | phenylpyrrole |
|  | metalaxyl-M 2.00% | phenylamide |
|  | thiabendazole 15.00% | benzimidazole |
|  | thiamethoxam 35.00% | insecticide |
| **Vibrance® Maxx** | fludioxonil 2.50% | phenylpyrrole |
|  | metalaxyl-M 3.70% | phenylamide |
|  | sedaxane 5.00% | carboxamide |
| **Acronis®** | pyraclostrobin 5.00% | strobilurin |
|  | thiophanate-methyl 45.00% | benzimidazole |
| **Vibrance® Cruiser Maxx** | metalaxyl-M 3.60% | phenylamide |
|  | fludioxonil 1.20% | phenylpyrrole |
|  | sedaxane 1.20% | carboxamide |
|  | thiamethoxam 24.00% | insecticide |
| **Topsin Flo** | thiophanate-methyl 50.00% | benzimidazole |
| **Topseed Extra** | thiophanate-methyl 15.00% | benzimidazole |
|  | ethaboxam 0.50% | thiazole carboxamide |
|  | thiram 30.00% | dithiocarbamate |

An 8-mm mycelial plug from a 7-day-old *T. koningiopsis* colony was placed at the center of each plate. Four replicates per concentration were incubated at 26 ± 2°C under continuous light for 10 days. The colony diameter was measured in two perpendicular directions, and the diameter of the mycelial plug was subtracted before calculating the mean diameter of the colony (MD). For each concentration/fungicide, mycelial growth inhibition (MGI), was calculated by MGI = ((MDc - MDi) / MDc) x 100, where MDc = mean colony diameter for the control (no fungicide added), and MDi = mean colony diameter of *T. koningiopsis* grown on medium added of fungicide. For each replicate of each fungicide concentration, MGIi values were linearly regressed on the logarithm (log_10_) of fungicide concentration to estimate the dose that inhibited mycelial growth by 50% (EC_50_ value). For each fungicide, the sensitivity was established according to criteria defined by Edgington et al. (1971): Highly fungitoxic: EC_50_ < 1 ppm; Moderately fungitoxic: 1–50 ppm; Non-fungitoxic: > 50 ppm.

### 2.3 Inoculum production

For *T. koningiopsis* inoculum, 125 g of sterile sorghum was inoculated with five 8 mm mycelium discs of the selected *Trichoderma* and incubated at 26 ± 2°C with continuous light for 10 days. *Trichoderma* conidia were washed from the sorghum grains with sterile distilled water and Tween 20 (0.1% v/v). Conidia concentration was determined by a Neubauer chamber.

For Mp inoculum, sterile rice was inoculated with Mp17 and incubated for 20 days at 30 ± 2°C in darkness to promote microsclerotia development.

### 2.4 Fields trials

Field trials were conducted in San Agustín, Cruz Alta department, Tucumán, Argentina (26°49’24.2“S 64°51’19.7”W), a field with a history of charcoal rot, during three soybean growing seasons (2021/2022, 2023/2024, and 2024/2025). The experimental design was a completely randomized block with four replicates. Each treatment consisted of four 3-m rows, spaced 0.5 m apart, and manually planted at a density of 23 seeds per meter.

Seed treatments included *T. koningiopsis* alone or combined with Acronis® or Topseed Extra (Table 2), previously selected in earlier *in vitro* assays. Chemical products were applied at label rates, and *Trichoderma* was applied at a concentration of 1×10^9^ conidia Tr/100 kg seed. When combined with Topseed Extra, the Tr dose was increased by 50%.

**Table 2.**
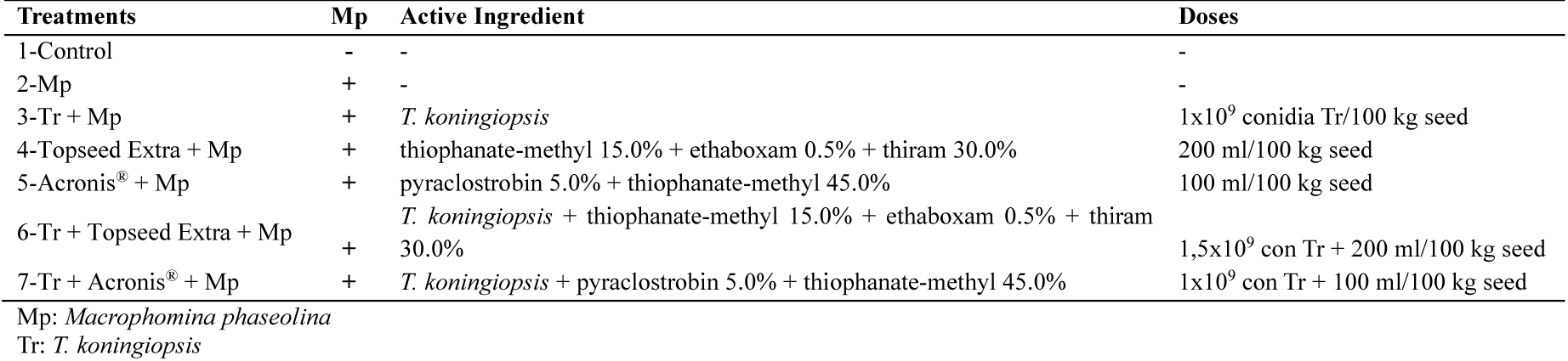
Seed treatments performed for *Macrophomina phaseolina* managment at field trial. San Agustín, Cruz Alta, Tucumán, Argentina. 2021/2022, 2023/2024 and 2024/2025 seasons.

| Treatments | Mp | Active Ingredient | Doses |
| --- | --- | --- | --- |
| 1-Control | - | - | - |
| 2-Mp | + | - | - |
| 3-Tr + Mp | + | *T. koningiopsis* | 1x10<sup>9</sup> conidia Tr/100 kg seed |
| 4-Topseed Extra + Mp | + | thiophanate-methyl 15.0% + ethaboxam 0.5% + thiram 30.0% | 200 ml/100 kg seed |
| 5-Acronis® + Mp | + | pyraclostrobin 5.0% + thiophanate-methyl 45.0% | 100 ml/100 kg seed |
| 6-Tr + Topseed Extra + Mp | + | *T. koningiopsis* + thiophanate-methyl 15.0% + ethaboxam 0.5% + thiram 30.0% | 1,5x10<sup>9</sup> con Tr + 200 ml/100 kg seed |
| 7-Tr + Acronis® + Mp | + | *T. koningiopsis* + pyraclostrobin 5.0% + thiophanate-methyl 45.0% | 1x10<sup>9</sup> con Tr + 100 ml/100 kg seed |

Mp: *Macrophomina phaseolina*
Tr: *T. koningiopsis*

To test the effectiveness of these treatments, at planting time, 10 g of colonized rice with Mp per linear meter was manually applied to each row. The trials were established on 11 January 2022, 09 January 2024 and 16 December 2024 for the respective growing seasons.

Plant emergence, disease severity, grain yield and yield increase respect to Mp control were evaluated. For emergence, the number of plants at 7, 14 and 21 days after planting (dap) were registered. Charcoal rot severity was recorded at stage R7 (Fehr and Caviness, 1977) using the scale of Paris et al. (2006) (1 = no discoloration and no microsclerotia visible; 2 = no discoloration of vascular tissue, with very few microsclerotia visible in the pith, vascular tissue or under the epidermis; 3 = partially discolored vascular tissue, with microsclerotia partially covering the tissue; 4 = discolored vascular tissue, with numerous microsclerotia visible in the tissue under the outer epidermis, in stem and root sections; 5 = vascular tissue with numerous microsclerotia producing a dark color inside and outside of the stem and root tissue). The samples were obtained by cutting 10 cm above and below soil level, including root and stem tissue, then washed and rinsed three times with tap water to remove any attached soil. Finally, crop yield (kg/ha) was measured at harvest time and yield increase respect to Mp control was calculated.

### 2.5 Environmental conditions data

Environmental conditions during the growing seasons 2021/2022, 2023/2024 and 2024/2025, were obtained from a nearby weather station located in San Agustín, Cruz Alta, Tucumán (26°49’24.2“S 64°51’19.7”W) from planting date to 21 dap and for all the soybean cycle. The following parameters were considered: rainfall (mm), number of days with rainfall, number of days with air temperatures >35°C and number of days with air temperatures >40°C (data from Sección - Agrometeorología EEAOC).

### 2.6 Effect of the temperature on *T. koningiopsis* mycelial growth

Mycelial growth was evaluated at 26°C, 28°C, 30°C, 32°C, 35°C, 40°C and 45°C on APDA. In Petri dishes of 9 cm, containing APDA (pH= 6), 8 mm mycelium discs of the bio-controller were placed in the center of the Petri dish. The plates were sealed with plastic film to avoid contamination and incubated at different temperatures during 92 h. Four replicated were used per each temperature evaluated. Colony diameter was measured at multiple time points using a digital caliper (Marathon Brand, Ontario, Canada) and the area under the curve (AUC) was calculated for each temperature.

### 2.7 Statistical analysis

The parameters percentage of plant emergence at 7, 14, and 21 dap, severity at R7, and soybean yield (kg/ha) were analyzed using a linear mixed-effects model including fixed effects of treatment, growing season, and their interaction, and a block effect. Heteroscedasticity among growing seasons was modeled using a VarIdent variance structure. Mean separation was performed using the DGC test (α = 0.05) in InfoStat (Di Rienzo et al., 2020), and model fitting was conducted in Navure (Navure, 2023). The parameter AUC was statistically evaluated with mixed linear models and a means comparison test (Tukey, α = 0.05) in InfoStat.

## 3. Results

### 3.1 *In vitro* compatibility of *T. koningiopsis* with chemical seed treatment fungicides

The results of the *in vitro* compatibility are presented in Table 3. Chemical fungicides Apron**^®^** Maxx RFC (metalaxyl-M 3.75% + fludioxonil 2.50%), Cruiser**^®^** Plus (fludioxonil 2.50% + difenoconazole 2.50% + thiamethoxam 26.30%), Cruiser**^®^** Advance (fludioxonil 2.50% + metalaxyl-M 2.00% + thiabendazole 15.00% + thiamethoxam 35.00%), Vibrance**^®^** Maxx (fludioxonil 2.50% + metalaxyl-M 3.75% + sedaxane 5.00%) and Topseed Extra (thiophanate-methyl 15.00% + ethaboxam 0.50% + thiram 30.00%) where classified as moderately fungitoxic.

**Table 3.** *In vitro* compatibility of chemical commercial fungicides against *T. koningiopsis*: EC_50_ values and sensitivity classification according to Edgington et al. (1971). Phytopathology Laboratory - EEAOC - Tucumán, Argentina.

| Commercial Fungicide | Active Ingredient(s) | EC<sub>50</sub><br>(ppm) | Sensitivity (Edgington et al., 1971) |
| --- | --- | --- | --- |
| **Apron® Maxx RFC** | metalaxyl-M 3.75% + fludioxonil 2.50% | 7.78 | Moderately fungitoxic |
| **Cruiser® Plus** | fludioxonil 2.50% + difenoconazole 2.50% + thiamethoxam 26.30% | 10.02 | Moderately fungitoxic |
| **Cruiser® Advance** | fludioxonil 2.50% + metalaxyl-M 2.00% + thiabendazole 15.00% + thiamethoxam 35.00% | 18.70 | Moderately fungitoxic |
| **Vibrance® Maxx** | fludioxonil 2.50% + metalaxyl-M 3.75% + sedaxane 5.00% | 26.19 | Moderately fungitoxic |
| **Acronis®** | pyraclostrobin 5.00% + thiophanate-methyl 45.00% | 60.77 | Non-fungitoxic |
| **Vibrance® Cruiser Maxx** | metalaxyl-M 3.60% + fludioxonil 1.20% + sedaxane 1.20% + thiamethoxam 24.00% | 104.80 | Non-fungitoxic |
| **Topsin Flo** | thiophanate-methyl 50.00% | 389.24 | Non-fungitoxic |
| **Topseed Extra** | thiophanate-methyl 15.00% + ethaboxam 0.50% + thiram 30.00% | 20.25 | Moderately fungitoxic |

On the other hand, Acronis^®^ (pyraclostrobin 5.00% + thiophanate-methyl 45.00%), Vibrance**^®^** Cruiser Maxx (metalaxyl-M 3.60% + fludioxonil 1.20% + sedaxane 1.20% + thiamethoxam 24.00%) and Topsin Flo (thiophanate-methyl 50.00%) registered EC_50_ values > 50 ppm; classified as non-fungitoxic under the Edgington scale.

For field trials, one chemical fungicide classified as non-fungitoxic (Acronis^®^) and another classified as moderately fungitoxic (Topseed Extra) were selected. When Topseed Extra was applied combined with *T. koningiopsis*, the dose of the biocontrol agent was increased by 50%.

### 3.2 Fields trials

The percentage of plant emergence at 7, 14, and 21 days after planting (dap) during the 2021/2022, 2023/2024, and 2024/2025 crop seasons are presented in Table 4. Charcoal rot disease was reproduced under field conditions using isolate Mp17, as evidenced by the reduction in percentage of plant emergence observed in treatment 2 (Mp inoculated control). Statistical differences were found in the first two seasons, whereas in 2024/2025 no significant differences were observed.

**Table 4.** Percentage of plant emergence at 7, 14 and 21 days after planting (dap), in field trials. San Agustín, Cruz Alta, Tucumán, Argentina. 2021/2022, 2023/2024 and 2024/2025 seasons.

| Treatments | 2021/2022 |  |  |  |  |  |  |  |  | 2023/2024 |  |  |  |  |  |  |  |  | 2024/2025 |  |  |  |  |  |
| --- | --- | --- | --- | --- | --- | --- | --- | --- | --- | --- | --- | --- | --- | --- | --- | --- | --- | --- | --- | --- | --- | --- | --- | --- |
|  | E. 7 dap |  |  | E. 14 dap |  |  | E. 21 dap |  |  | E. 7 dap |  |  | E. 14 dap |  |  | E. 21 dap |  |  | E. 7 dap |  | E. 14 dap |  | E. 21 dap |  |
|  | Mean | SE |  | Mean | SE |  | Mean | SE |  | Mean | SE |  | Mean | SE |  | Mean | SE |  | Mean | SE | Mean | SE | Mean | SE |
| 1-Control | 72.4 | 7.9 | A | 75.8 | 7 | A | 69.1 | 6.2 | A | 89.9 | 2.4 | B | 86.7 | 2.55 | B | 86.4 | 2.6 | B | 39.7 | 7.1 | 37.9 | 8.0 | 36.2 | 8.1 |
| 2- Mp | 35.2 | 7.9 | B | 35.7 | 7 | B | 33.0 | 6.2 | B | 76.1 | 2.4 | C | 77.8 | 2.55 | B | 73.7 | 2.6 | C | 25.8 | 7.1 | 21.4 | 8.0 | 20.3 | 8.1 |
| 3- Tr + Mp | 29.4 | 7.9 | B | 30.1 | 7 | B | 28.3 | 6.2 | B | 86.0 | 2.4 | B | 86.6 | 2.55 | B | 84.5 | 2.6 | B | 32.9 | 7.1 | 28.6 | 8.0 | 28.0 | 8.1 |
| 4- TopSeed Extra + Mp | 70.1 | 7.9 | A | 70.7 | 7 | A | 63.4 | 6.2 | A | 95.4 | 2.4 | A | 94.8 | 2.55 | A | 93.1 | 2.6 | A | 30.2 | 7.1 | 28.6 | 8.0 | 40.1 | 8.1 |
| 5- Acronis® + Mp | 65.9 | 7.9 | A | 79.9 | 7 | A | 74.1 | 6.2 | A | 96.3 | 2.4 | A | 91.3 | 2.55 | A | 91.8 | 2.6 | A | 57.2 | 7.1 | 55.2 | 8.0 | 53.4 | 8.1 |
| 6- Tr + TopSeed Extra + Mp | 52.1 | 7.9 | A | 57.9 | 7 | A | 54.7 | 6.2 | A | 84.9 | 2.4 | B | 82.0 | 2.55 | B | 81.0 | 2.6 | B | 40.5 | 7.1 | 39.5 | 8.0 | 50.4 | 8.1 |
| 7- Tr + Acronis® + Mp | 76.0 | 7.9 | A | 85.0 | 7 | A | 80.9 | 6.2 | A | 87.6 | 2.4 | B | 82.4 | 2.55 | B | 80.8 | 2.6 | B | 44.4 | 7.1 | 45.6 | 8.0 | 46.1 | 8.1 |
| P value | 0.0013 |  |  | 0.0001 |  |  | <0.0001 |  |  | 0.0003 |  |  | 0.0029 |  |  | 0.0008 |  |  | 0.0899 |  | 0.1010 |  | 0.0696 |  |
| df | 18 |  |  | 18 |  |  | 18 |  |  | 18 |  |  | 18 |  |  | 18 |  |  | 18 |  | 18 |  | 18 |  |

SE: Standard error df: degrees of freedom Pairwise means comparison method: DGC, alpha=0.05 Means with a common letter are not significantly different (p>0.05)

During the 2021/2022 growing season, percentage of plant emergence differed statistically at 7, 14, and 21 dap (*p* ≤ 0.0013). The non-inoculated control showed significantly higher emergence than the Mp inoculated control at all evaluation dates. The Mp and Tr + Mp treatments showed the lowest emergence values at all evaluation dates. In contrast, all other seed treatments presented significantly higher emergence than Mp control.

In the 2023/2024 growing season, significant differences in percentage of plant emergence were observed between treatments at all evaluation dates (*p* ≤ 0.0029). As in the previous season, the non-inoculated control showed significantly higher emergence than the Mp inoculated control at 7 and 21 dap. The Mp inoculated control produced the lowest values at 7, 14, and 21 dap. At 7 and 21 dap, all the treatments showed significantly higher plant emergence than the Mp inoculated control. The treatments that showed significantly higher plant emergence were TopSeed Extra + Mp and Acronis® + Mp, followed by Control, Tr + Mp, Tr + TopSeed Extra + Mp and Tr + Acronis® + Mp. In this season, the combined treatments (Tr + TopSeed Extra + Mp and Tr + Acronis® + Mp) showed lower emergence values than the chemical treatments (TopSeed Extra + Mp and Acronis® + Mp) at all evaluation dates.

During the 2024/2025 growing season, although all seed treatments showed higher plant emergence values compared to the Mp control, no significant differences among treatments were found in this parameter at 7, 14, and 21 dap (*p* ≥ 0.0696).

Over the growing seasons, the interaction percentage of plant emergence × growing season was statistically significant (*p* < 0.0001) (Table 5). Mean comparisons showed statistical difference among the three seasons. The 2023/2024 crop season presented the highest percentage of plant emergence (86.2%), followed by 2021/2022 (59.0%) and 2024/2025 (38.2%). These results indicate that environmental conditions varied over years, with 2023/2024 growing season providing the most favorable conditions for plant emergence, while 2024/2025 growing season showed the lowest emergence percentages.

**Table 5.** Average of plant emergence at 2021/2022, 2023/2024 and 2024/2025 growing seasons in field trials. San Agustín, Cruz Alta, Tucumán, Argentina.

| Growing season | Average of emergencies | SE |  |
| --- | --- | --- | --- |
| 2021/2022 | 59.0 | 1.4 | B |
| 2023/2024 | 86.2 | 0.6 | A |
| 2024/2025 | 38.2 | 1.6 | C |
| *P value* | <0.0001 |  |  |
| *df* | 228 |  |  |

SE: Standard errors df: degrees of freedom Pairwise means comparison method: DGC, alpha=0.05. Means with a common letter are not significantly different (p>0.05).

The growing seasons also interacted with treatment responses (growing seasons × treatment, *p* < 0.0001), demonstrating that treatment performance depended on the growing season conditions as shown in Table 6. Although, within each season, the Mp inoculated control exhibited the lowest emergence value, confirming the negative impact of inoculation on early stand establishment (Table 7). Chemical seed treatments such as Acronis® + Mp and TopSeed Extra + Mp generally achieved the highest emergence values. In the 2023/2024 growing season, Acronis® + Mp and TopSeed Extra + Mp treatments presented significantly higher values compared to both controls. *Trichoderma*-based treatments showed intermediate performance, improving emergence compared to Mp inoculated control but with lower values than the only chemical treatments. In 2024/2025 growing season, the percentage of plant emergence was lower in all treatments, and no significant difference was detected between the inoculated and non-inoculated control.

**Table 6.** Average plant emergence across growing seasons for different treatments in field trials. San Agustín, Cruz Alta, Tucumán, Argentina.

| Treatments | Average of emergencies | SE |  |
| --- | --- | --- | --- |
| 1-Control | 66.0 | 1.9 | B |
| 2- Mp | 44.3 | 1.9 | C |
| 3- Tr + Mp | 48.3 | 1.9 | C |
| 4- TopSeed Extra + Mp | 65.1 | 1.9 | B |
| 5- Acronis® + Mp | 73.9 | 1.9 | A |
| 6- Tr + TopSeed Extra + Mp | 60.3 | 1.9 | B |
| 7- Tr + Acronis® + Mp | 69.9 | 1.9 | A |
| *P value* |  | <0.0001 |  |
| df |  | 228 |  |

SE: Standard errors
df: degrees of freedom
Pairwise means comparison method: DGC, alpha=0.05.
Means with a common letter are not significantly different ( $p>0.05$ ).

**Table 7.** Year interaction with treatment responses in plant emergence across growing seasons in field trials. San Agustín, Cruz Alta, Tucumán, Argentina.

| Treatments | Growing season | Average of emergencies | SE |  |
| --- | --- | --- | --- | --- |
| 1-Control | 2021/2022 | 72.4 | 3.7 | C |
| 2- Mp |  | 34.6 | 3.7 | E |
| 3- Tr + Mp |  | 29.3 | 3.7 | E |
| 4- TopSeed Extra + Mp |  | 68.0 | 3.7 | C |
| 5- Acronis® + Mp |  | 73.3 | 3.7 | C |
| 6- Tr + TopSeed Extra + Mp |  | 54.9 | 3.7 | D |
| 7- Tr + Acronis®+ Mp |  | 80.6 | 3.7 | B |
| 1-Control | 2023/2024 | 87.7 | 1.5 | B |
| 2- Mp |  | 75.9 | 1.5 | C |
| 3- Tr + Mp |  | 85.7 | 1.5 | B |
| 4- TopSeed Extra + Mp |  | 94.4 | 1.5 | A |
| 5- Acronis® + Mp |  | 93.2 | 1.5 | A |
| 6- Tr + TopSeed Extra + Mp |  | 82.6 | 1.5 | B |
| 7- Tr + Acronis® + Mp |  | 83.6 | 1.5 | B |
| 1-Control | 2024/2025 | 37.9 | 4.1 | E |
| 2- Mp |  | 22.5 | 4.1 | E |
| 3- Tr + Mp |  | 29.9 | 4.1 | E |
| 4- TopSeed Extra + Mp |  | 33.0 | 4.1 | E |
| 5- Acronis® + Mp |  | 55.3 | 4.1 | D |
| 6- Tr + TopSeed Extra + Mp |  | 43.5 | 4.1 | D |
| 7- Tr + Acronis® + Mp |  | 45.4 | 4.1 | D |
| *P value* |  | <0.0001 |  |  |
| *df* |  | 228 |  |  |

SE: Standard errors
df: degrees of freedom
Pairwise means comparison method: DGC, alpha=0.05.
Means with a common letter are not significantly different (p>0.05).

The reduction in percentage of plant emergence across growing seasons may be attributed to the environmental conditions recorded during the evaluated periods (Table 8). The lowest percentage of emergence recorded in the 2024/2025 growing season may have been associated with limited precipitations during the assessment period (19.77 mm) and 12 days with temperatures >35°C. In the 2021/2022 season, 191.98 mm of precipitations were recorded, however 11 days >35°C and 6 days >40°C were registered that could have affected soybean emergency. On the other hand, the 2023/2024 growing season, which showed the highest emergence values, registered 74.18 mm of precipitations and only 4 days with temperatures >35°C.

**Table 8.** Precipitation (mm), number of days with precipitation, number of days with air temperature >35°C and number of days with air temperature >40°C. From planting time to 21 dap. San Agustín, Cruz Alta, Tucumán, Argentina. 2021/2022, 2023/2024 and 2024/2025 seasons.

| Season | Evaluated period | Precipitation (mm) | Days with precipitation | Days with T >35°C | Days with T >40°C |
| --- | --- | --- | --- | --- | --- |
| 2021/2022 | Planting time to 7dap | 50.79 | 3 | 7 | 6 |
|  | 7 dap to 14 dap | 57.39 | 2 | 2 | 0 |
|  | 14 dap to 21 dap | 83.80 | 3 | 2 | 0 |
|  | **Total** | **191.98** | **8** | **11** | **6** |
| 2023/2024 | Planting time to 7dap | 25.15 | 5 | 1 | 0 |
|  | 7 dap to 14 dap | 49.03 | 2 | 1 | 0 |
|  | 14 dap to 21 dap | 0.00 | 0 | 2 | 0 |
|  | **Total** | **74.18** | **7** | **4** | **0** |
| 2024/2025 | Planting time to 7dap | 0.51 | 1 | 5 | 0 |
|  | 7 dap to 14 dap | 2.03 | 2 | 3 | 0 |
|  | 14 dap to 21 dap | 17.23 | 3 | 4 | 0 |
|  | **Total** | **19.77** | **6** | **12** | **0** |

Root severity evaluated at the R7 growth stage did not differ significantly among treatments in any of the evaluated seasons (*p* ≥ 0.0735) (Table 9). In 2021/2022, severity values were low among treatments (2.0 to 2.6), but no significant difference was found between the non-inoculated and the Mp inoculated control. During the 2023/2024 season, severity values range from 2.2 to 3.0, without statistical differences among treatments. In 2024/2025, disease severity was higher (2.7 to 3.3) which corresponded to hot and dry conditions (Figure 1).

**Figure 1.**
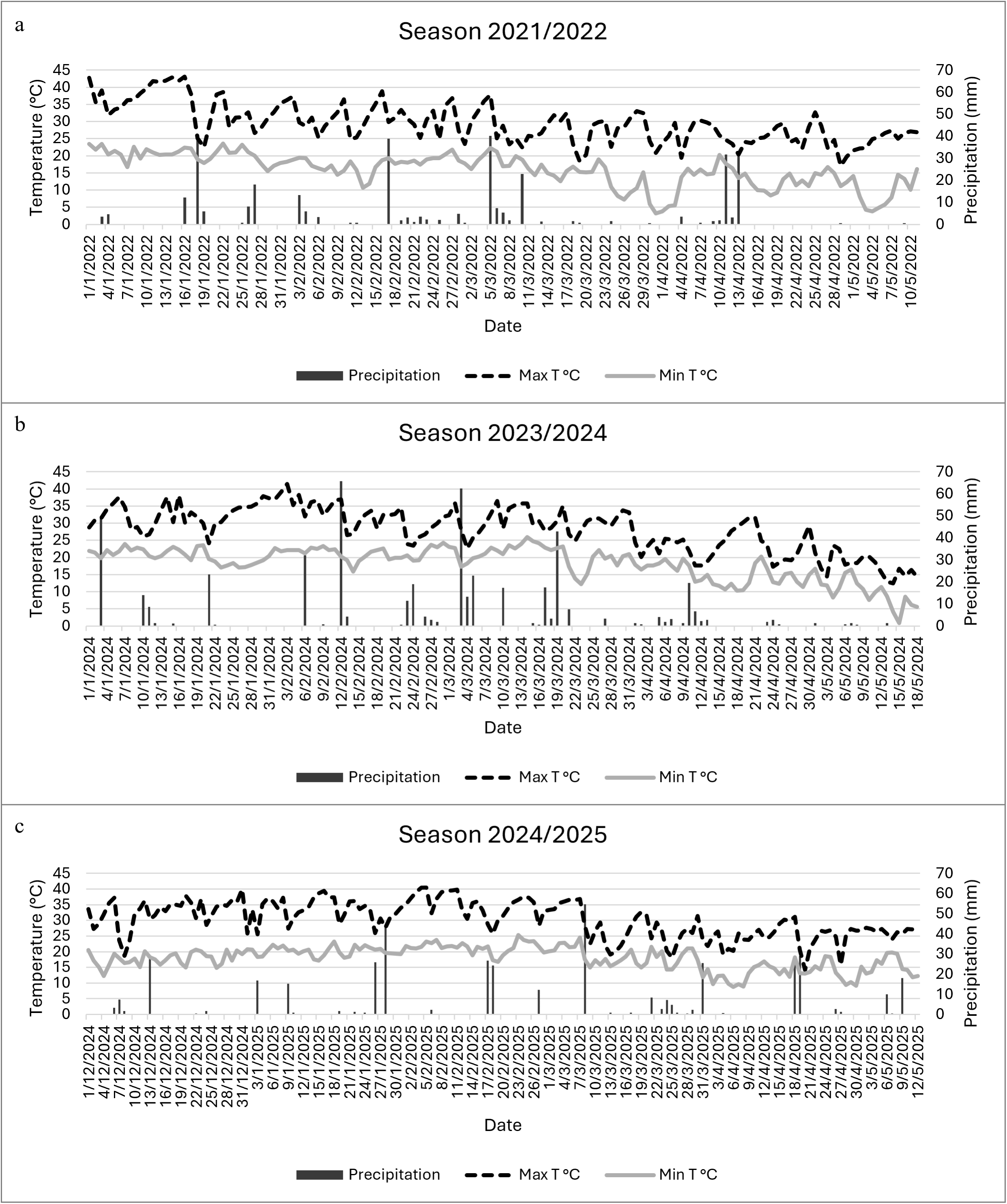
Daily data for precipitation (mm), maximum and minimum air temperature (◦C). San Agustín, Cruz Alta, Tucumán, Argentina. **a-**2021/2022, **b-**2023/2024 and **c-**2024/2025 seasons.

**Table 9.** Charcoal rot severity values on a scale from 1 to 5 in field trials. San Agustín, Cruz Alta, Tucumán, Argentina. 2021/2022, 2023/2024 and 2024/2025 seasons.

| Treatments | 2021/2022 |  | 2023/2024 |  | 2024/2025 |  |
| --- | --- | --- | --- | --- | --- | --- |
|  | Mean | SE | Mean | SE | Mean | SE |
| 1-Control | 2.3 | 0.3 | 2.7 | 0.3 | 1.7 | 0.4 |
| 2- Mp | 2.6 | 0.3 | 3.0 | 0.2 | 2.8 | 0.5 |
| 3- Tr + Mp | 2.4 | 0.3 | 2.5 | 0.2 | 2.7 | 0.3 |
| 4- TopSeed Extra + Mp | 2.3 | 0.3 | 2.2 | 0.2 | 2.8 | 0.3 |
| 5- Acronis® + Mp | 2.3 | 0.3 | 2.3 | 0.2 | 4.0 | 0.5 |
| 6- Tr + TopSeed Extra + Mp | 2.0 | 0.3 | 2.4 | 0.2 | 2.7 | 0.3 |
| 7- Tr + Acronis® + Mp | 2.0 | 0.3 | 2.8 | 0.2 | 3.3 | 0.3 |
| *P value* | 0.6854 |  | 0.1467 |  | 0.0735 |  |
| *df* | 17 |  | 17 |  | 13 |  |

SE: Standard errors
df: degrees of freedom
Pairwise means comparison method: DGC, alpha=0.05.
Means with a common letter are not significantly different (p>0.05).

The interaction root severity × growing season was statistically significant (*p* = 0.0056) (Table 10). Mean comparisons showed statistical difference among the three seasons. The 2021/2022 crop season presented the lowest root severity values (2.3) with statistical differences respect to the 2023/2024 (2.6) and the 2024/2025 (2.8) growing seasons.

**Table 10.** Average of charcoal rot severity at 2021/2022, 2023/2024 and 2024/2025 growing seasons in field trials. San Agustín, Cruz Alta, Tucumán, Argentina.

| Growing season | Average of charcoal rot severity | SE |  |
| --- | --- | --- | --- |
| 2021/2022 | 2.3 | 0.1 | B |
| 2023/2024 | 2.6 | 0.1 | A |
| 2024/2025 | 2.8 | 0.2 | A |
| *P value* | 0.0056 |  |  |
| *df* | 53 |  |  |

SE: Standard errors df: degrees of freedom Pairwise means comparison method: DGC, alpha=0.05. Means with a common letter are not significantly different (p>0.05).

The severity of charcoal rot at the R7 stage was associated with the environmental conditions registered in each soybean growing season. During the 2021/2022 season, high maximum temperatures (often >35°C) and irregular rainfall distribution were favorable for charcoal rot development resulting in moderate mean disease severity values (2.3) (Figure 1a).

In contrast, the 2023/2024 season showed higher mean disease severity (2.6), due to less precipitation and frequent drought stresses registered during the reproductive stages of soybean (Figure 1b). The 2024/2025 season was characterized by persistent high maximum temperatures (frequently >38°C) and prolonged dry periods with few rainfall events. Such conditions are optimal for charcoal rot development and correspond to the highest mean disease severity observed (2.8).

Grain yield did not differ significantly among treatments in any of the evaluated seasons (*p* ≥ 0.1043) (Table 11). In 2021/2022, yields were relatively low across treatments, ranging from 741.7 to 1070.8 kg/ha, with no significant differences; however, several seed treatments showed yield increases of 200.0 to 329.2 kg/ha compared to the Mp inoculated control. In 2023/2024, yields were higher (3737.8 to 4375.6 kg/ha), indicating favorable environmental conditions, and although yield increases were obtained with the seed treatments, the statistical analysis revealed no significant differences. Finally, in the 2024/2025 growing season, no significant differences were detected among treatments, however, chemical seed treatments and the combined seed treatments showed higher yields respect to both controls.

**Table 11.** Mean yield (kg/ha) and mean yield increase respect to Mp control (kg/ha) in field trials. San Agustín, Cruz Alta, Tucumán, Argentina. 2021/2022, 2023/2024 and 24/2025 seasons.

| Treatments | 2021/2022 |  |  | 2023/2024 |  |  | 2024/2025 |  |  |
| --- | --- | --- | --- | --- | --- | --- | --- | --- | --- |
|  | Yield (kg/ha) | SE | Yield increase (kg/ha) | Yield (kg/ha) | SE | Yield increase (kg/ha) | Yield (kg/ha) | SE | Yield increase (kg/ha) |
| 1-Control | 1020.8 | 187.6 | 279.2 | 3745.8 | 180.4 | 46.5 | 1163.9 | 272.7 | -145.3 |
| 2- Mp | 741.7 | 187.6 |  | 3737.8 | 208.1 |  | 1291.7 | 313.6 |  |
| 3- Tr + Mp | 941.7 | 187.6 | 200.0 | 4375.6 | 180.4 | 637.8 | 1108.6 | 313.6 | -183.1 |
| 4- TopSeed Extra + Mp | 1062.5 | 187.6 | 320.8 | 4316.3 | 180.4 | 578.5 | 1474.2 | 272.7 | 182.5 |
| 5- Acronis® + Mp | 1070.8 | 187.6 | 329.2 | 4153.0 | 180.4 | 415.2 | 1559.7 | 272.7 | 268.0 |
| 6- Tr + TopSedd Extra + Mp | 1058.3 | 187.6 | 316.6 | 4141.1 | 180.4 | 403.3 | 1621.7 | 272.7 | 330.0 |
| 7- Tr + Acronis® + Mp | 1000.0 | 187.6 | 258.3 | 4262.0 | 180.4 | 524.2 | 1577.4 | 272.7 | 285.8 |
| *P value* | 0.8607 |  |  | 0.1043 |  |  | 0.7561 |  |  |
| df | 18 |  |  | 17 |  |  | 16 |  |  |

SE: Standard errors df: degrees of freedom Pairwise means comparison method: DGC, alpha=0.05 Means with a common letter are not significantly different (p>0.05)

The effect of growing season on yield was statistically significant (*p* < 0.0001) (Table 12), while neither treatment (*p* = 0.2035) or growing season × treatment interaction (*p* = 0.8291) was significant (data not shown). Mean comparisons of yields showed statistical difference among growing seasons, 2023/2024 presented the highest yields (4100.1 kg/ha), followed by 2024/2025 (1401.7 kg/ha) and 2021/2022 (985.1 kg/ha). The absence of significant growing season × treatment interaction indicates that treatment effect on yield did not vary across seasons and that yield differences were affected by environmental conditions.

**Table 12.** Average of yield at 2021/2022, 2023/2024 and 2024/2025 growing seasons in field trials. San Agustín, Cruz Alta, Tucumán, Argentina.

| Growing season | Average yield (kg/ha) | SE |  |
| --- | --- | --- | --- |
| 2021/2022 | 985.1 | 74.6 | C |
| 2023/2024 | 4100.1 | 67.7 | A |
| 2024/2025 | 1401.7 | 102.0 | B |
| *P value* | <0.0001 |  |  |
| *df* | 57 |  |  |

SE: Standard errors df: degrees of freedom Pairwise means comparison method: DGC, alpha=0.05. Means with a common letter are not significantly different (p>0.05).

The differences in yield among seasons may be explained by the contrasting rainfall regimes. The 2021/2022 growing season accumulated only 334.74 mm of rainfall, below the optimal requirement (550.00 to 700.00 mm) for soybean in northwestern Argentina. The pronounced water deficit, especially during reproductive stages, limited yield potential. In 2023/2024, total rainfall increased to 440.15 mm, with better distribution throughout the crop cycle, resulting in the highest yields. In 2024/2025, the combination of precipitations below the critical threshold for optimal yield (372.08 mm), heat and drought stress during pod filling led to low productivity.

Seed treatments that included only *T. koningiopsis* did not differ statistically from the Mp control in all of the parameters evaluated during the 2021/2022 and 2024/2025 seasons. The occurrence of several days with air temperature >35°C (17 days in 2021/2022 and 12 days in 2024/2025 season) could have affected negatively *T. koningiopsis*. For this reason, the growth of *T. koningiopsis* was evaluated under different temperatures conditions.

### 3.3 Effect of the temperature on the mycelial growth of *Trichoderma koningiopsis*

The *in vitro* mycelial growth of *T. koningiopsis* was evaluated across a temperature gradient ranging from 26°C to 45°C. Mycelial growth at different times is shown at Figure 2 and the area under the curve (AUC) is presented in Table 13.

**Figure 2.**
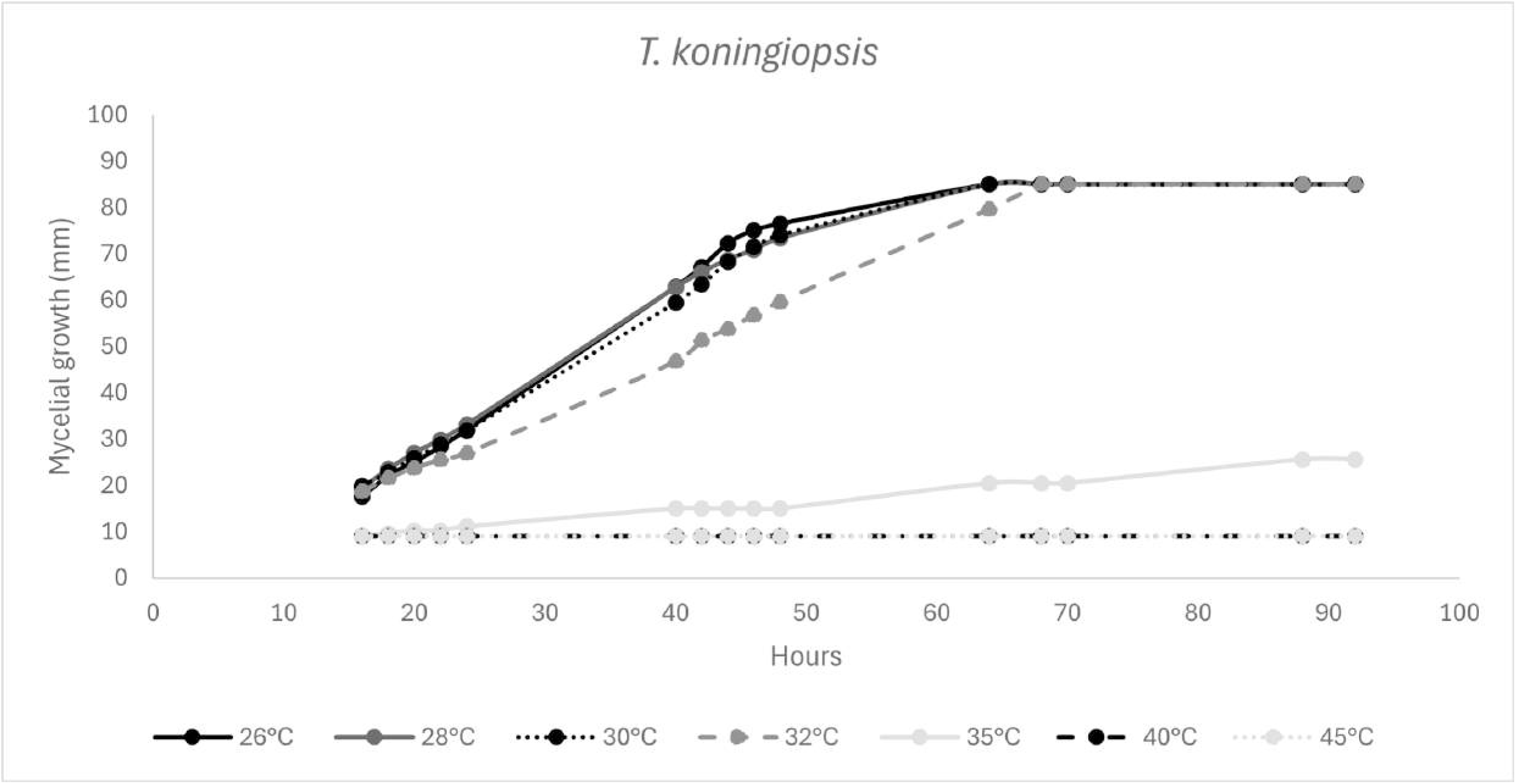
Growth curves of *T. koningiopsis* at different temperatures. Phytopathology Laboratory – EEAOC - Tucumán, Argentina.

**Table 13.** Area under the curve (AUC) for *T. koningiopsis* growth at different temperatures. Phytopathology Laboratory - EEAOC - Tucumán, Argentina.

| Temperature | AUC | SE |  |
| --- | --- | --- | --- |
| 26 °C | 4928.5 | 108.4 | C |
| 28 °C | 5172.4 | 108.4 | C |
| 30 °C | 5129.6 | 108.4 | C |
| 32 °C | 4682.5 | 108.4 | C |
| 35 °C | 1334.0 | 108.4 | B |
| 40 °C | 666.0 | 108.4 | A |
| 45 °C | 666.0 | 108.4 | A |
| *P value* |  | <0,0001 |  |
| *df* |  | 18 |  |

SE: Standard errors
df: Degrees of freedom
Pairwise means comparison method: Tukey $\alpha=0.05$
Means with a common letter are not significantly different ( $p>0.05$ )

The optimal mycelial growth of *T. koningiopsis* was observed at 26°C to 30°C. Growth declined noticeably at 32°C to 35°C and was absent at 40°C to 45°C, indicating strong thermal inhibition or potential loss of fungal viability. Plates previously incubated at 40°C and 45°C during 60 h, were subsequently incubated at 26°C and no growth was observed, confirming irreversible thermal damage.

These findings confirm that *T. koningiopsis* is thermally sensitive, with an optimal growth range between 28°C to 32°C, and a critical threshold for inhibition above 35°C.

The AUC parameter presented statistical differences (*p* < 0.0001) at different temperatures evaluated. The highest AUC values were observed at 28°C and 30°C, indicating optimal conditions for mycelial growth, followed by AUC at 26°C and 32°C with no significant differences. At 35°C, AUC declined and temperatures above resulted in a reduction and minimal AUC values detected at 40°C and 45°C.

## 4. Discussion

The present study evaluated the compatibility of *T*. *koningiopsis* with commercial fungicides and its biocontrol effect, alone or combined with chemical seed treatments, for managing charcoal rot in soybean across three contrasting growing seasons. The results demonstrated that treatment responses were strongly modulated by environmental conditions, particularly temperature and water availability, which affected both disease development and the efficacy of biological control. Our multi-season study revealed that environmental restrictions, particularly high temperatures, were the dominant factor of biological seed treatment performance.

The *in vitro* compatibility assays showed differences in fungitoxicity among the chemical commercial fungicides evaluated. The results ranged from moderately fungitoxic to non-fungitoxic according to the Edgington et al. (1971) scale. These findings are consistent with previous studies reported by Manandhar et al. (2020) who evaluated *in vitro* compatibility of five *Trichoderma* isolates with different chemical fungicides. They showed that Bavistin (carbendazim 50%), Cryzole (hexaconazole 5%), Benlate (benomyl 50%) and Saaf (carbendazim 12% + mancozeb 63%) inhibited *Trichoderma* growth. On the other hand, *Trichoderma* isolates were compatible with Krilaxyl (metalaxyl 8% + mancozeb 64%) and Aver up (chlorothalonil 75%).

In the field trials performed over three growing seasons, the Mp-inoculated control consistently exhibited the lowest emergence, confirming the negative effect of *M. phaseolina* on early stand establishment. Most seed treatments improved emergence relative to the inoculated control, but the statistical significance varied among years. The highest mean emergence occurred in the 2023/2024 growing season, characterized by moderate temperatures and adequate rainfall, whereas 2024/2025 growing season showed the lowest mean emergence under severe heat and drought stress conditions.

The significant growing season effect and the treatment × growing season interaction detected indicate that treatment performance was not stable across seasons. The results also highlight that chemical seed treatments tended to present better percentage of emergences than biological or combined treatments under unfavorable conditions, suggesting that *Trichoderma* activity may have been restricted in the hottest and driest environments. This is consistent with the thermal sensitivity observed *in vitro* where mycelial growth declined above 32 °C and was completely inhibited at 40 to 45 °C, with irreversible loss of viability. These temperatures were exceeded several times during the emergence period, particularly in the 2021/2022 and 2024/2025 growing seasons, indicating that *T. koningiopsis* may have lost viability before it was able to establish and persist on soybean roots. Under these conditions, the biocontrol agent likely failed to colonize the rhizosphere, which would explain its reduced capacity to mitigate the effects of *M. phaseolina* in the field.

Similar results were described by Cavalcante et al. (2025). They evaluated three *Trichoderma* isolates, two *T. asperellum* and one *T. longibrachiatum*, growth at 5 to 45°C. The optimal temperature described was 27°C and no mycelial growth was observed at 45°C. However, the isolates showed growth when they were re-incubated at 27°C, indicating that the temperature had not lethal effects on those isolates. This response emphasizes the importance of species-specific thermal tolerance when selecting biocontrol agents for environments where charcoal rot is prevalent.

Despite reductions in severity values in some treatments, no statistically significant differences were detected in any season. On the other hand, the interaction of root severity × growing season was statistically significant, associated with the environmental conditions registered of each growing season. Zaccaron et al. (2024) evaluated the effect of drought stress during soybean development on charcoal rot. They observed higher severity values when irrigation was interrupted at the R5 stage in Mp-inoculated plots compared with the irrigated plots throughout the entire crop cycle. These results demonstrate that predisposing environmental conditions are required for charcoal rot development.

Yield differences among treatments were not statistically significant in any of the seasons evaluated, although chemical seed treatments and the combined seed treatments showed higher yields respect to both controls. Zandoná et al. (2019) reported yield increases of up to 14% when soybean seeds were treated with fludioxonil combined with *Trichoderma* spp., highlighting that certain chemical–biological combinations can enhance crop performance under field conditions. Their findings indicate that fludioxonil-based formulations are compatible with *Trichoderma* spp. and may contribute to improved plant development and grain yield. However, in the present study, such benefits were not consistently expressed across seasons, likely due to the strong influence of environmental stress on treatment performance.

Although growing season significantly affected yield, no significant growing season × treatment interaction was detected. These results indicate that treatment effects on yield did not vary across seasons, and that yield variation was mainly explained by environmental conditions.

The differences registered in percentage of plant emergence between treatments did not correspond with the yield results observed in this study. This may be explained by the soybean plant’s remarkable capacity to compensate for low plant populations through the development of axillary branches that produce additional pods and take advantage of open space. These branches allow individual plants to occupy gaps in the stand, and such plants can yield two to three times more than plants growing under normal population densities (Suhre et al., 2014).

## 5. Conclusions

Our findings indicate that seed treatments with *T. koningiopsis* improve the percentage of plant emergencies under favorable environmental conditions in plots with *M. phaseolina* inoculations. It was also demonstrated that this *Trichoderma* is compatible with certain chemical fungicides. However, their performance in the field was strongly influenced by temperature, and their contribution to disease management was limited under severe heat and drought stress. These results highlight the importance of selecting biocontrol strains with thermal tolerance and integrating biological and chemical tools within management strategies that consider environmental conditions.

## Acknowledgments

We would like to thank Dr. Andrea Natalia Peña Malavera (ITANOA) for performing part of the statistical analyses included in this study. This work was supported by grants from EEAOC and CONICET (Argentina).

## Data availability

The data that support the findings of this study are available from the corresponding author upon reasonable request.

